# Bulk sample testing by real-time PCR for detecting *Diaporthe destruens*-infected seedlings of sweet potatoes

**DOI:** 10.1101/2025.03.11.642721

**Authors:** Takashi Fujikawa, Hiroe Hatomi, Kohji Yamamura, Yoshifumi Tsukiori, Yasuhiro Inoue, Kazuki Fujiwara

## Abstract

Foot rot disease of sweet potato is a fungal disease that causes serious damage to sweet potato production. It is especially important to check the health of seedlings as there is a risk that if the seed tubers or stem seedlings infected with foot rot pathogen are introduced into the production fields, it will rapidly spread throughout the fields. We have already developed a real-time PCR method to diagnose sweet potatoes contaminated with foot rot pathogen. In this study, to apply this method to the bulk sample testing, we provided examples on estimating the “probability of detection (POD)” for detecting bulk samples, and determining the number of subgroups and the number of samples in each subgroup for the bulk sample testing. Our results indicate that, for tubers, when testing between 100 and 1200 fragments per bulk sample, the lowest detection rate was approximately 81% at 800 fragments. For stems, when testing between 100 and 600 fragments, the lowest detection rate was approximately 83% at 500 fragments. Based on the estimated POD, the required sample size for bulk testing was calculated as follows: for the impermissible threshold proportion of contamination of infected plants (p) of 1%, between 390 and 624 samples are needed, while for p of 0.1%, between 3,840 and 6,228 samples are required. The bulk sample testing method using real-time PCR demonstrated in this study will become a powerful quality testing method for checking the health of sweet potato seedlings, and will make a significant contribution to sweet potato production.

## Introduction

Foot rot disease of sweet potato (*Ipomoea batatas* (L.) Lam.) is caused by a fungus *Diaporthe destruens* (Harter) Hirooka, Minosh. & Rossman (Gai et al. 2016; Maeda et al. 2022). Infected sweet potatoes show blackening and wilting of the stem (foot rot), followed by the death of the entire plant. In infected root tubers, root rot is often developed during storage. Severely affected fields frequently also have black, hardened stem lesions and rotting tubers in the soil (Gai et al. 2016; Kobayashi 2019; Maeda et al. 2022; Nishioka et al. 2021). The disease was first reported in the United States in 1912 and has since been found in Cuba, the Caribbean, Brazil, Argentina, eastern Africa, New Zealand, and other countries (Fujiwara et al. 2021; Gai et al. 2016; Harter 1913). Recently, it has also spread to East Asia, including Japan, Taiwan, China, and Korea, and is becoming a major problem for sweet potato production (Gai et al. 2016; Huang et al. 2012; Paul et al. 2019). In particular, since the planting of infected seedlings (stem seedlings or seed tubers) is the most primary infection source (Usui and Kushima 2020), securing healthy seedlings is important for controlling this disease and preventing its spread. The cultivation of sweet potatoes as an agricultural product begins with the planting of stem seedlings in the fields. These stem seedlings are prepared and shipped through (1) the multiplication of seed tubers, the production and multiplication of stem seedlings derived from seed tubers, and (2) the multiplication of stem seedlings derived from *in vitro* culture. During these seedlings production process, abnormalities including symptom of foot rot disease are visually removed. However, it is difficult to identify seedlings that are latently or slightly infected with foot rot. Therefore, we developed a genetic detection method using real-time PCR that can identify foot rot pathogen contamination without relying on visual inspection (Fujiwara et al. 2021; Tsukiori et al. 2023). The real-time PCR method developed by Fujiwara et al. (2021) for detecting sweet potato foot rot and dry rot fungi is capable of identifying even a small amount of pathogen DNA. This method can be used to diagnose infections by two similar pathogens in abnormal tubers and stems found in cultivation fields and so on. Furthermore, based on this method, we developed endogenous control primers for sweet potato to confirm the success of DNA extraction, and examined the process from DNA extraction to PCR analysis to update the diagnostic method for foot rot disease (Tsukiori et al. 2023). These methods made it possible to diagnose foot rot disease without relying on visual inspection, but they involved examining each individual tuberous root or stem, and were not methods for testing many samples at once. Thus, to detect the presence of foot rot-infected sweet potato samples from bulk sweet potato samples, we performed a statistical study on the number of samples that should be taken when inspecting the health of seedlings during the seedling production process.

## Materials and Methods

### Samples of sweet potato plants

Seed tubers and stem seedlings of sweet potato (The varieties were mostly “Kogane Sengan”) were collected from various fields in areas where control measures were insufficient and foot rot disease was widespread at the start of this study. Among these, samples that were confirmed in advance to be infected with *Diaporthe destruens* were extracted and used as naturally derived positive samples in this study. In addition, commercially available or cultivated tubers and stems (The varieties were mainly “Kogane Sengan”, but sometimes included other varieties such as “Beni Azuma”.) were used as healthy samples. For processing, tubers and stems were washed with distilled water, and their surfaces were sterilized by wiping with the cleaning tissue Kimwipe papers (Nippon Paper Crecia Co., Ltd., Tokyo, Japan) soaked with 70% ethanol before cutting. Followingly, the tubers were cut into about 1-cm^3^ pieces, and the stems were cut into small pieces of about 1.5-cm length. The samples were either used immediately or frozen until use. Frozen samples were thawed in a warm bath at 37°C for 30 min.

### DNA extraction

When performing the bulk sample testing, it is necessary to grind several hundred samples at once and extract DNA entirely. To make this possible, we decided to crush the samples using a stomacher machine, which is used for food and seed testing, and extract DNA from the homogenate solutions. The schematic diagram is shown in Figure 1.

**Figure 1.**
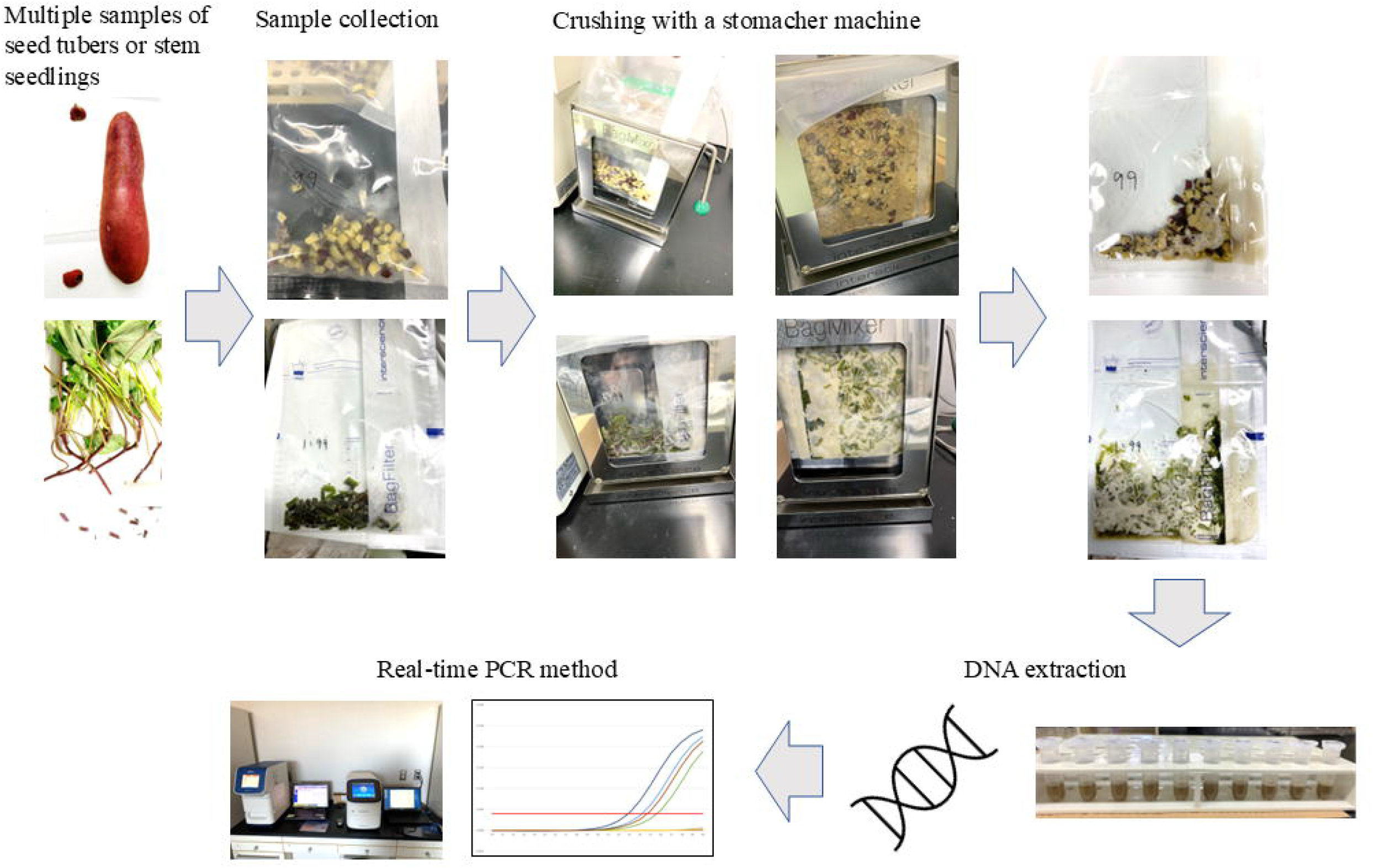
Process diagram for crushing multiple samples, DNA extraction, and real-time PCR. The entire process of sample crushing, DNA extraction, and real-time PCR for the multiple sample testing is illustrated. The samples were crushed using a stomacher machine, which enabled sufficient DNA extraction.

Respective bulk tuber blocks or stem fragments including each positive sample were put into the sterile stomacher bags with filter, Bagpage (Interscience, France) and AP1 buffer of a DNeasy plant mini kit were added to the bags at the adequate amounts (6-ml/100-200 tuber blocks, 15-ml/400-600 tuber blocks, 18-ml/800 tuber blocks, 20-ml/1000-1200 tuber blocks, 6-ml/100 stems, and 15-ml/400-600 stems). The samples in Bagpage were crushed in a stomacher machine Bag mixer (Interscience) for 5 min, then the bags were turned over and crushed for an additional 5 min. By this treatment, most of the samples were crushed, deformed, and powdered, and the AP1 buffer became liquids in which the sweet potato components were dissolved. Then, from behind the filter, each extract was dispensed in replicates of 600 μL into 1.5 ml tubes. In the subsequent steps, DNA was extracted using a DNeasy plant mini kit according to the manufacturer’s instructions.

### Real-time PCR

Real-time PCR was performed with slight modifications to the method described by Fujiwara et al. (2021) and Tsukiori et al. (2023) as follows: A QuantStudio 3 real-time PCR system (Thermo Fisher Scientific Inc., MA) with KOD SYBR qPCR Mix (Toyobo Co., Ltd., Osaka, Japan) was used according to the manufacturer’s protocol for real-time PCR. The 20-μl reaction mixtures contained 10 μl of qPCR Mix, 0.04 μl of 50× ROX reference dye, 0.5 μl of the respective 10 μM primer sets (forward and reverse) with 2 μl of template DNA (10-fold diluted extracted DNA). For detection of foot rot disease pathogen, *D. destruens*, the forward primer Dd ITS-F (5’-GTTTTTATAGTGTATCTCTGAGC-3’) and the reverse primer Dd ITS-R (5’-GGCCTGCCCCCTTAAAAA-3’) (Fujiwara et al. 2021), and for detection of sweet potato internal control, the forward primer Ipo psaB-F (5’-TGTGAAACGTTACCCTGCCA-3’) and the reverse primer Ipo psaB-R (5’-GGACCCGGAGACTTTTTGGT-3’) (Tsukiori et al. 2023) were used respectively. The PCR protocol consisted of initial denaturation at 94 □ for 1 min, and 40 cycles of denaturation at 96 □ for 30 s, annealing at 55 □ for 30 s, and extension at 72 □ for 30 s. Further denaturation at 96 □ for 15 s, holding at 55 □ for 1 min, and heating from 55 to 96 □ for15 s were carried out for melting curve analysis. We have previously reported that the real-time PCR method used in this study can be used without falsely identifying other pathogens as positive (Tsukiori et al. 2023) and used as a detection method with a low false-negative rate (Fujikawa et al. 2025) by setting the cutoff Ct value at 35; therefore, the results in this test were also judged using a cutoff Ct value of 35.

### Probit regression and determination of “the probability of detection”

When bulk specimens including a positive (*D. destruens-*contaminated) sample are tested together in real-time PCR assay, the target DNA (*D. destruens* ITS region) may be diluted, resulting in a negative result. To perform the bulk sample testing without missing infectious sample contamination, it is necessary to understand “the probability of inspection errors” in real-time PCR assay for the bulk sample testing. “The probability of inspection errors” is synonymous with “the probability of non-detection”, which is calculated as “1 - the probability of detection (POD)”. According to Yamamura et al. (2019), under the assumption that the probability of detection curve (POD curve) monotonically increases as the proportion of infected samples in multiple samples increases, the POD value may be obtained from the estimated POD curve or results (observed values) detected in each bulk sample testing. To evaluate the success of bulk sample testing in tuber samples, DNA was extracted and real-time PCR was performed from the following bulk groups: 100 (99 healthy + 1 infected), 200 (199 healthy + 1 infected), 400 (399 healthy + 1 infected), 600 (599 healthy + 1 infected), 800 (799 healthy + 1 infected), 1000 (999 healthy + 1 infected), and 1200 (1199 healthy + 1 infected). Similarly, to evaluate the success of bulk sample testing in stem samples, DNA was extracted and real-time PCR was performed from the following bulk groups: 100 (99 healthy + 1 infected), 400 (399 healthy + 1 infected), 500 (499 healthy + 1 infected), and 600 (599 healthy + 1 infected). Infected samples used in this study were fragments of seed tubers or stem seedlings naturally infected with the foot rot pathogen in the field. For infected samples, real-time PCR assays were performed on 15 randomly collected samples of infected plants to confirm that the Ct value was low enough (clearly positive), and the 95% confidence interval was calculated for the mean Ct value. In each test, DNA extraction and real-time PCR were replicated 2 to 10 times from the same bulk sample. These tests were repeated a minimum of six times and a maximum of 16 times (Table 1). For each bulk sample, the observed detection rate was obtained from the number of successful detections in the number of repeated tests. Estimation of POD by probit regression and generation of POD curves were performed by the glm function in R4.2.1 (e.g., Suppl. Text 1).

**Table 1.** Changes in detection rate due to different in the number of pieces in samples.

a. Seed tubers
| **Number of pieces in group** | **Detection number of test sets (response)** | **Total number of test sets** | **Observed detection rate** | **Observed non-detection rate** |
| --- | --- | --- | --- | --- |
| **100** | **9** | **9** | **1** | **0** |
| **200** | **10** | **12** | **0.833333333** | **0.166666667** |
| **400** | **10** | **12** | **0.833333333** | **0.166666667** |
| **600** | **9** | **11** | **0.818181818** | **0.181818182** |
| **800** | **13** | **16** | **0.8125** | **0.1875** |
| **1000** | **7** | **7** | **1** | **0** |
| **1200** | **6** | **7** | **0.857142857** | **0.142857143** |

b. Stem seedlings
| **Number of pieces in group** | **Detection number of test sets (response)** | **Total number of test sets** | **Observed detection rate** | **Observed non-detection rate** |
| --- | --- | --- | --- | --- |
| **100** | **7** | **7** | **1** | **0** |
| **400** | **5** | **5** | **1** | **0** |
| **500** | **5** | **6** | **0.833333333** | **0.166666667** |
| **600** | **6** | **6** | **1** | **0** |

### Required sample size for the bulk sample testing

The minimum sample size to be tested must be determined in order to perform the bulk sample testing for health inspection on seed tubers or stem seedlings. If we can draw the sample completely at random from the entire lot, the required total sample size, *n*, can be calculated as follows (Hughes et al. 2002; Yamamura et al. 2016; 2019):

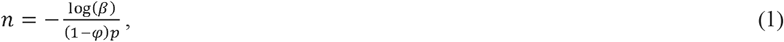

where *p* is the impermissible threshold proportion of infected seed tubers or stem seedlings, *β* is the consumer’s risk indicating the probability that an impermissible lot is mistakenly accepted, and *φ* is the probability of inspection error which is the value given by 1−POD. The value of 0.05 has been adopted for *β* in the Japanese plant quarantine inspection since 1992 (Yamamura et al. 2016). If only one infected tuber (or seedling) is included in the test portion from which the DNA is extracted, the concentration of pathogen tubers (or seedlings) is given by 1/*n*. The target DNA may be too diluted to be detected by real-time PCR if *n* is too large. Therefore, it is necessary to divide the *n* tubers (or seedlings) into *g* test portions for performing real-time PCR. Let *m* be the number of tubers (or seedlings) that are handled in one test portion (thus *n*= *gm*). Now, let *φ* be the maximum probability of non-detection when one or more infected tubers are included in these *m* tubers (or seedlings) that is when the concentration is not smaller than 1/*m* Yamamura et al. (2019) discussed a composite sampling, which is a kind of non-random sampling, and derived a conservative sample size. Then, they additionally suggested the required sample size for random sampling as a limit of composite sampling. The equation for required sample size, *n*, is as follows:

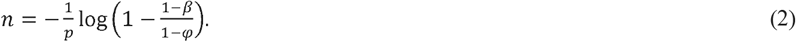

However, we can use much efficient formulae if we can perform a complete random sampling; we can replace *β* by *β* ^(1/*g*)^in equation (2), because we are performing independent *g* tests for controlling the consumers risk at *β*. Then, the required number of groups, *g*, should satisfy the following inequation:

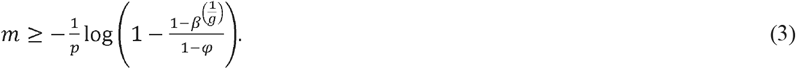

The total sample size, *n*, should satisfy the following inequation:

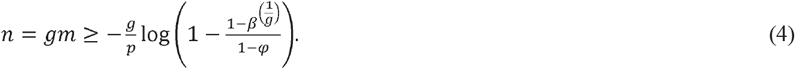

The number of groups *g* may be determined by finding the minimum *g* that satisfy inequation (3). For example, we can find the appropriate number of groups, *g*, by evaluating inequation (3) by changing *g* from *g* = 1 to 10. Suppl. Text 2 shows two example scripts that are performed using R4.2.1. Please notice that inequation (4) essentially reduces to equation (1) when *g* becomes large.

## Results

### Estimating “the probability of detection” for the bulk sample testing

As described in Materials and Methods, when testing bulk samples using real-time PCR, the target DNA may be diluted below the limit of detection by the DNA solution of healthy samples, resulting in negative results. When considering this possibility, it is necessary to estimate “the probability of detection (POD)” in the bulk sample testing. For the bulk sample testing of seed tubers, multiple plots consisting of 100, 200, 400, 600, 800, 1000, and 1200 tuber blocks were examined. We led each plot contained one infected tuber block. The infected fragments used here were derived from tubers that had actually become diseased in the field, and the Ct value was calculated using 15 blocks randomly selected from multiple infected fragments; the Ct value was 25.399 ± 2.224 with a 95% confidence interval (Suppl. Table 1a). Similarly, for the bulk sample testing of stem seedlings, multiple plots consisting of 100, 400, 500, and 600 stem fragments were examined. And we led each plot contained one infected stem fragment. The infected fragments used here were also derived from stems that had actually become diseased in the field, and the Ct value was calculated using 15 blocks randomly selected from multiple infected fragments; the Ct value was 25.504 ± 2.478 with a 95% confidence interval (Suppl. Table 1b). The average Ct values of both tuber blocks and stem fragments were approximately 25, which was a sufficiently low Ct value. Therefore, since there was a difference of about 10 cycles from the real-time PCR cutoff Ct value of 35, the positive result was expected clearly indicated. For seed tubers and stem seedlings, respective plots of the bulk sample testing were performed (Suppl. Table 2 and Suppl. Table 3), the detection rate was observed from the detected number of test sets in the total number of test sets in each plot (Table 1). For seed tubers, the detection rate was 100% when there were 100 tuber blocks per group. As the number of blocks per group increased (i.e., the amount of foot rot pathogen DNA became increasingly diluted), the detection rate decreased (approximately 83% in 200 and 400 blocks, approximately 82% in 600 blocks, and approximately 81% in 800 blocks) but returned to 100% detection rate at 1,000 blocks. On the other hand, for stem seedlings, the detection rate dropped to approximately 83% only when there were 500 stem fragments per group, but when there were between 100 and 600 fragments per group, the detection rate was 100%. In both cases, the lowest detection rate was approximately 81% when 800 blocks were included per group of seed tubers. Since the detection rate varied depending on the number of pieces in a group, we estimated “the probability of detection (POD)” using a probit regression. Here, the POD in one laboratory was calculated using R (without inter-laboratory testing). Probit regression curves for POD were generated for seed tubers and stem seedlings, respectively (Suppl. Fig 1). However, since the approximation formula were not sufficiently significant in both cases, it was inappropriate to set the limit of detection (LOD) from the estimated POD when each group contained 800 pieces (m = 800, 1/m = 0.00125). For this reason, we decided to use the lowest detection rate actually observed this time, the value at 800 pieces of seed tubers (detection rate = 0.8125), as a conservative POD estimate.

### Determination of the parameters that make valid the bulk sample testing

For the bulk sample testing method using the real-time PCR for detecting foot rot pathogen to be effective in investigating healthy seedlings, it is necessary to clarify how many randomly selected seedlings need to be tested in one test (lot). For this sampling lot testing, it is important to set “the inspection level” that indicates the permissible threshold proportion of contamination of infected plants, and “the consumer’s risk” that indicates the probability that a lot containing infected plants will be mistakenly determined to be healthy. In addition, the incidence rate of detection errors due to bulk samples is indicated using the POD value. By setting these values, it is possible to calculate the number of samples required for inspection, so that even with seed tubers or stem seedlings, which have an infinite lot size, a negative result (determined not to be infected with the foot rot pathogen) can be proven with a limited number of random samples. The required sample number can be calculated using equations described in “Materials and Methods.” The impermissible threshold proportion of contamination of infected plants (*p*) was set to 0.01 and 0.001 to suppose the introduction of seed tubers into disinfection processes and stem seedlings into fields. The consumer’s risk (*β*) was set to 0.05, which is the value generally used in plant quarantine. The probability of inspection error (*φ*) was “1-POD,” where POD was set to 0.8125, the observed value from the above test results, and *φ* was set to 0.1875. Consequently, the total sample number (*n*), as well as the number of subgroups (*g*) and the number of test samples within each subgroup (*m*) were calculated (Table 2). When the impermissible threshold proportion of contamination of infected plants (*p*) was set to 0.01, the total number of samples required for testing (*n*) ranged from 390 to 624 for the numbers of subgroups (*g*) between 2 and 10 (Table 2a). The number of test samples within each subgroup (*m*) was a maximum of 312 (*n*=624, *g* =2) and a minimum of 39 (*n*=390, *g* =10). Moreover, when the impermissible threshold proportion of contamination of infected plants (*p*) was set to 0.001, the total number of samples required for testing (*n*) ranged from 3840 to 6228 for the numbers of subgroups (*g*) between 2 and 10 (Table 2b). The number of test samples within each subgroup (*m*) was a maximum of 3114 (*n*=6228, *g* =2) and a minimum of 384 (*n*=3840, *g* =10). In the bulk sample detection method used in this study, a maximum of 1200 samples were used for seed tubers and a maximum of 600 samples for stem seedlings (Table 1). Therefore, if the maximum detection amount confirmed in this study is considered to be 1200 fragments for seed tubers and 600 fragments for stem seedlings, the conditions to satisfy this *m* are all cases of *g* = 2 to 10 when *p* = 0.01, and *g* ≥ 4 for seed tubers and *g* ≥ 7 for stem seedlings when *p* = 0.001.

**Table 2.**
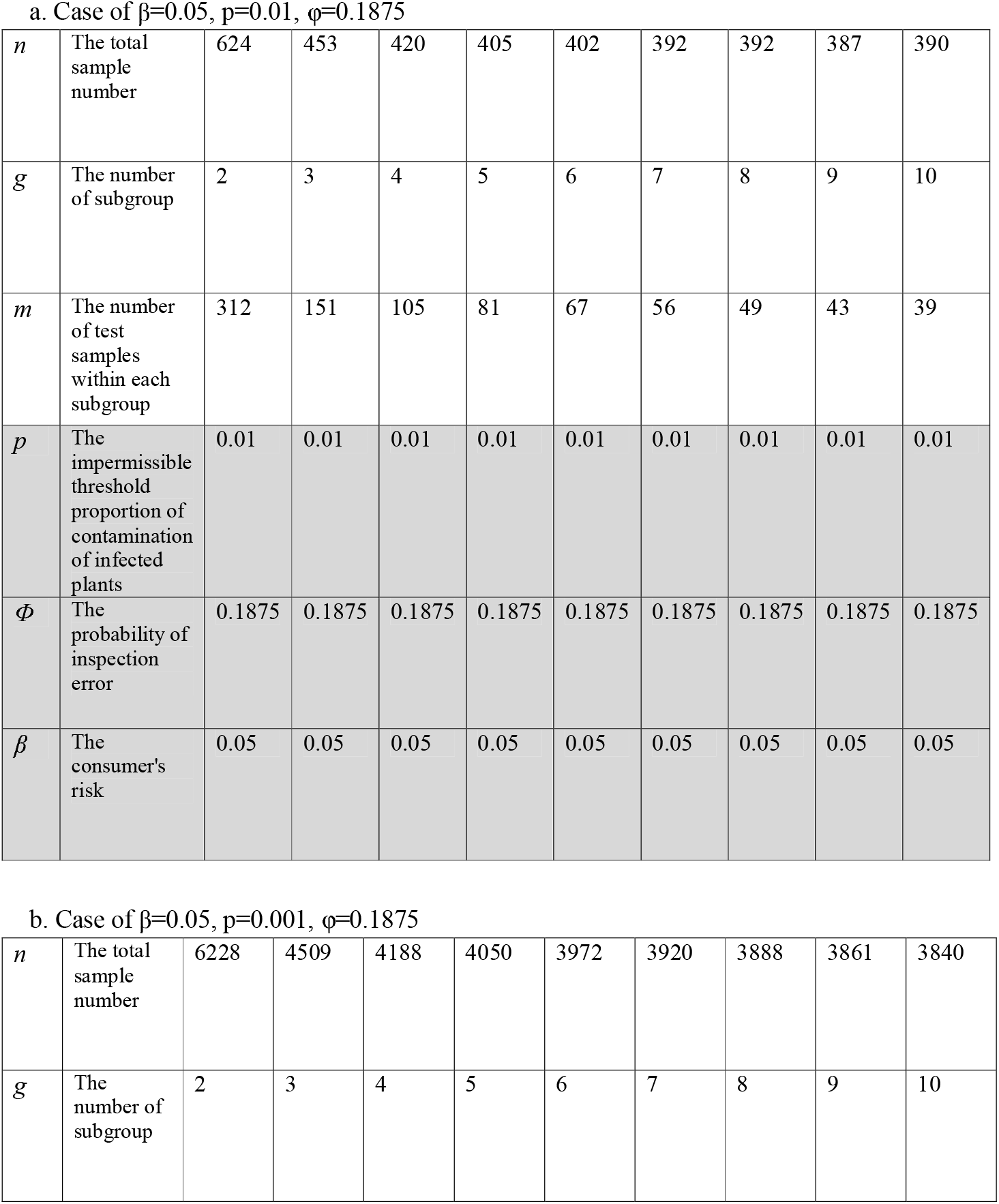
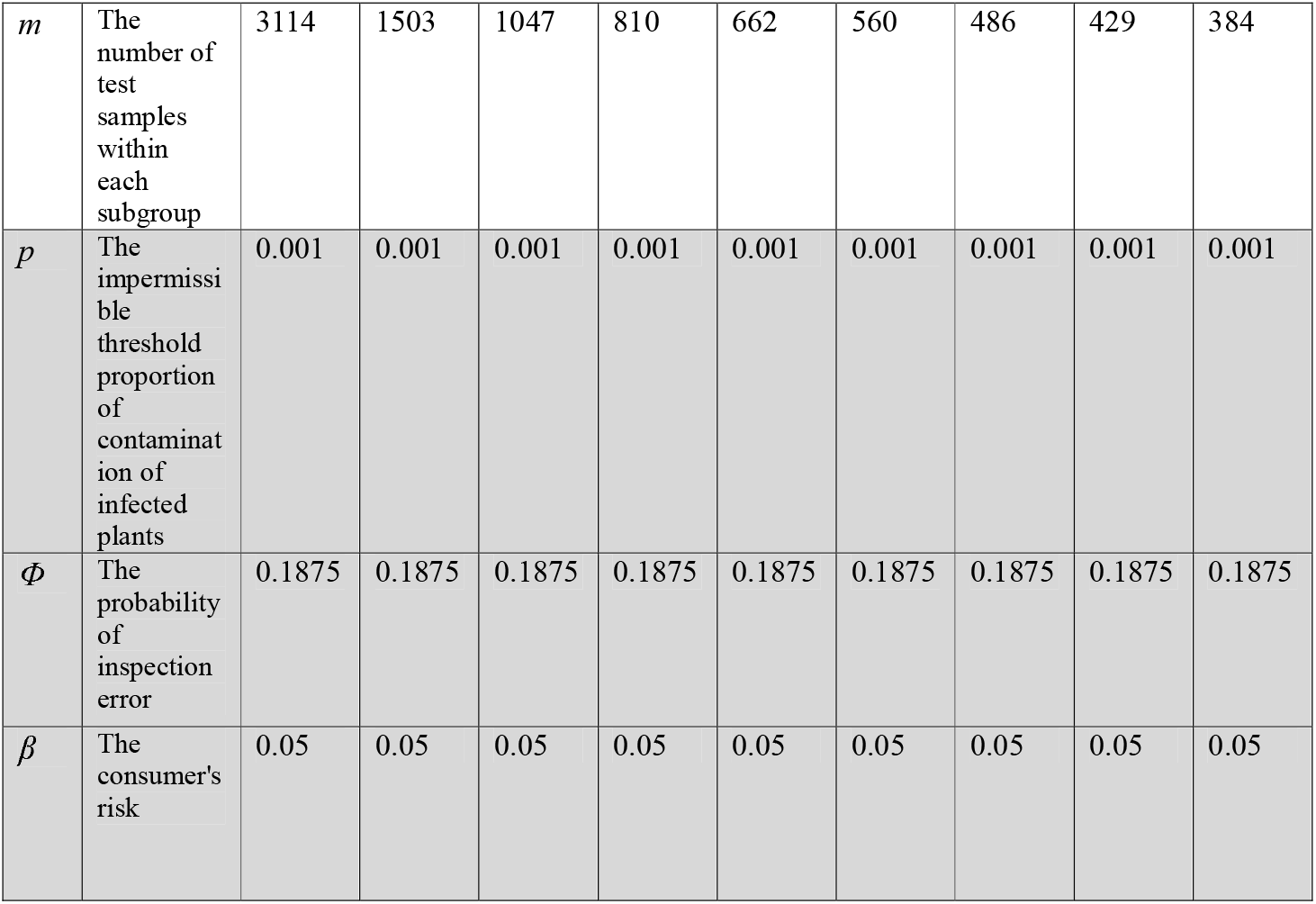
Determination of sampling manner for the bulk sample testing.

a. Case of $\beta=0.05$ , $p=0.01$ , $\phi=0.1875$
|  |  |  |  |  |  |  |  |  |  |  |
| --- | --- | --- | --- | --- | --- | --- | --- | --- | --- | --- |
| $n$ | The total sample number | 624 | 453 | 420 | 405 | 402 | 392 | 392 | 387 | 390 |
| $g$ | The number of subgroup | 2 | 3 | 4 | 5 | 6 | 7 | 8 | 9 | 10 |
| $m$ | The number of test samples within each subgroup | 312 | 151 | 105 | 81 | 67 | 56 | 49 | 43 | 39 |
| $p$ | The impermissible threshold proportion of contamination of infected plants | 0.01 | 0.01 | 0.01 | 0.01 | 0.01 | 0.01 | 0.01 | 0.01 | 0.01 |
| $\phi$ | The probability of inspection error | 0.1875 | 0.1875 | 0.1875 | 0.1875 | 0.1875 | 0.1875 | 0.1875 | 0.1875 | 0.1875 |
| $\beta$ | The consumer's risk | 0.05 | 0.05 | 0.05 | 0.05 | 0.05 | 0.05 | 0.05 | 0.05 | 0.05 |
| b. Case of $\beta=0.05$ , $p=0.001$ , $\phi=0.1875$ |  |  |  |  |  |  |  |  |  |  |
| $n$ | The total sample number | 6228 | 4509 | 4188 | 4050 | 3972 | 3920 | 3888 | 3861 | 3840 |
| $g$ | The number of subgroup | 2 | 3 | 4 | 5 | 6 | 7 | 8 | 9 | 10 |
| $m$ | The number of test samples within each subgroup | 3114 | 1503 | 1047 | 810 | 662 | 560 | 486 | 429 | 384 |
| $p$ | The impermissible threshold proportion of contamination of infected plants | 0.001 | 0.001 | 0.001 | 0.001 | 0.001 | 0.001 | 0.001 | 0.001 | 0.001 |
| $\Phi$ | The probability of inspection error | 0.1875 | 0.1875 | 0.1875 | 0.1875 | 0.1875 | 0.1875 | 0.1875 | 0.1875 | 0.1875 |
| $\beta$ | The consumer's risk | 0.05 | 0.05 | 0.05 | 0.05 | 0.05 | 0.05 | 0.05 | 0.05 | 0.05 |

## Discussion

The real-time PCR method for detecting foot rot pathogen of sweet potato developed by Fujiwara et al. (2021) can detect the pathogen with high sensitivity even in tubers and seedlings, which are difficult to inspect visually. Thereby, this method is becoming widely used for diagnosing sweet potatoes in Japan. As well, detection techniques including PCR, have been widely developed in the field of plant protection. However, most of them are for diagnostic or identification purposes, specifically detecting a single suspect sample or an isolated pathogen, and there are not many reported methods available for testing bulk samples. In sweet potato production, it is important not only to diagnose foot rot disease in the field, but also to monitor for the contamination of foot rot pathogen during seedling production and supply. Therefore, there was a demand for adaptation to the bulk sample testing using the real-time PCR. When using the real-time PCR method to test the healthiness of sweet potato seedlings, it is not realistic to test each individual seed tuber and stem seedling. Generally, bulk samples are tested at the same time. Here, we considered that the bulk sample testing could be performed using the real-time PCR method of Fujiwara et al. (2021). In fact, experimental results confirmed that detection was possible even if one positive sample fragment was included among several hundred or more healthy sample fragments. In addition, we addressed the question of how many samples could be used to apply the bulk sample testing in the healthiness test of seedlings in actual inspection site. The “actual inspection site” referred to here is the site where the healthiness of seedlings is inspected when seed tubers possibly contaminated with foot rot pathogen are subjected to a disinfection process such as vapor heat treatment, or when disinfected seed tubers or cloned stem seedlings propagated in seedlings production facilities are brought to cultivation fields. At actual inspection sites, some parameters varies depending on the seedling production process and the purpose of the inspection, so it was determined based on interviews with agricultural corporations and various related organizations regarding their inspection needs. As already known from the report by Yamamura and the colleagues (Hughes et al. 2002; Yamamura et al. 2016; 2019), by carrying out appropriate sampling on a lot consisting of a large number of samples (theoretically up to an infinite size), it is possible to confirm the healthiness of the entire original lot. In order to clarify the number of total samples at this time, it is necessary to determine the parameters *p, β*, and *φ* based on the above equations in “Materials and Methods.” As described in above “Results”, the impermissible threshold proportion of contamination of infected plants (*p*) was set to 0.01 and 0.001 to suppose the introduction of seed tubers into disinfection processes and stem seedlings into fields. When introducing seed tubers or stem seedlings into fields, it is desirable that all seedlings are healthy, but it is necessary to mathematically determine the maximum percentage of contamination that is acceptable. In Japan, several thousand of sweet potato seedlings are planted per 10 ares (for the brewing variety Kogane Sengan, 3,000 seedlings are planted per 10 ares). Based on interviews with agricultural corporations and related parties, it was estimated that farmers seem to be able to visually detect and remove around three abnormal plants among approximately 3,000 seedlings planted in a 10-are field during the cultivation period. In other words, even if the seedlings are contaminated with foot rot pathogen at ca. 0.1%, it is believed that farmers can prevent the spread of the disease during cultivation. Furthermore, various control measures for use in fields have now been developed and improved (NARO et al. 2023; Nomiyama 2023; Shima et al. 2024), and it is expected that even the introduction of seedlings with a contamination rate of 0.1% will significantly suppress the spread of the disease. Therefore, considering the risk of introducing contaminated seedlings into fields, we set the impermissible threshold proportion of contamination of infected plants (*p*) to 0.001. Additionally, in disinfection processes for sweet potato seed tubers, vapor heat treatment is increasingly being used as a disinfection method. Vapor heat treatment is a heat treatment using saturated steam, which can directly kill pests through heat and is used for fruit import quarantine and so on. In fact, it is used for disinfection of sweet potato stem tubers to kill the sweet potato weevil, *Cylas formicarius*, the west Indian sweet potato weevil, *Euscepes postfasciatus*, and the sweet potato vine borer, *Omphisa anastomosalis* (Shimabukuro et al. 1997). This disinfection method is highly effective against foot rot disease, and experiments have shown that a 100-minute treatment at a constant temperature of 48°C and a constant 95% relative humidity can disinfect the foot rot pathogen while minimizing the impact on seed tubers (NARO 2023). Then, of 210 tubers that had been artificially inoculated with foot rot pathogen, the fungus was not re-isolated in 202 of them after vapor heat treatment, so it is thought that the disinfection effect was over 90 percent. When seedlings with a contamination rate of 0.1% are permissible to be introduced into the field as mentioned above, even if the contamination rate of the seed tubers before vaper heat treatment is 1%, it is expected that the contamination rate will decrease to 0.1% or less after vaper heat treatment. Therefore, we set the impermissible threshold proportion of contamination of infected plants (*p*) for tests of healthiness before seed tubers disinfection to 0.01. The consumer’s risk (*β*) was set to 0.05, a value commonly used in plant quarantine, to maintain realistic validity. When considering implementation in the bulk sample testing, the probability of inspection error (*φ*) was the most difficult to set. Although high accuracy can be achieved when diagnosing each individual sample using real-time PCR, there is a risk that when testing bulk samples, the target DNA may be diluted below the limit of detection, resulting in the false negative result. Considering this risk, we calculated the POD and estimated the probability of inspection error (*φ*) as 1-POD. This real-time PCR for detecting *D. destruens* was found to be quite robust even when used with bulk samples, and was capable of detecting a contaminated fragment from both tuber fragments and stem fragments in large numbers. As this result in this study, we felt that it would be more appropriate to use actual observed values for POD rather than estimating them using probit regression. Surely, PODs can vary between inspection sites, laboratories, and even between test operators. In some cases, probit regression estimates may be more valid than observed values. When actually testing bulk samples, the test operators must set up this POD carefully. To comprehensively set the POD, inter-laboratory testing at multiple inspection sites or laboratories is essential, and probit regression estimates can be applied as the POD. Therefore, if a comprehensive POD for this method is required, the inter-laboratory testing will be necessary as the future works. Depending on the impermissible threshold proportion of contamination of infected plants (*p*), the total number of samples required for healthiness testing of the lot (*n*), the number of subgroups (*g*), and the number of test samples within each subgroup (*m*) were calculated. Considering the practicality of the bulk sample testing, it is better to have a small number of subgroups (*g*), but this increases the number of test samples within each subgroup (*m*) as shown in Table 2. In the seedlings healthiness test, it is important to determine *g* and *m*, and it is necessary to know in advance the number of pieces in (sub)group that can be detected without any problems. In this study, the conditions for efficient bulk sample testing are considered to be from *g* = 3 (*m* = 151 at this time) to *g* = 5 (*m* = 81 at this time) when *p* = 0.01, and from *g* = 7 (*m* = 560 at this time) to *g* = 10 (*m* = 384 at this time) when *p* = 0.001. Here, the former is assumed the healthiness test of seed tubers, where *n* and *m* are the number of tubers and the test uses two pieces from one tuber (the neck and the tail). the pathogen is expected to have invaded through the stem and into the neck of tubers in the field. However, it may be difficult to distinguish the neck and tail in harvested tubers, so it is thought that both the neck and the tail should be examined. Thus, note that when *g* = 3, one subgroup actually contains 302 tuber pieces, and when *g* = 5, one subgroup contains 162 tuber pieces. The latter is assumed the healthiness test of stem seedlings, where *n* and *m* are the number of seedlings, and it is sufficient to test the ends of that number of seedlings. In order to ensure the health of sweet potatoes, we have developed a bulk sample testing method using real-time PCR to distinguish seed tubers and stem seedlings infected with foot rot pathogen. This method was developed by analyzing them by combining mathematical science and genetic experiments so that they can be used for bulk samples. In the future, this method is expected to become a powerful testing method for checking the health of sweet potato seedlings, which will greatly contribute to sweet potato production.

## Acknowledgements

We thank the members of the Institute for Plant Protection, NARO and the Kyushu Okinawa Agricultural Research Center, NARO for helpful discussions.

## Author contributions

TF, KY, YT, YI, and KF conceived and designed the study. TF, HH performed the experiments. TF, and KY contributed to the data analysis. TF, KY, YT, YI, and KF contributed to the manuscript writing.

## Ethical approval

This article does not contain any studies with human participants or animals.

## Data availability statement

The authors confirm that all data supporting the findings of this study are available within the article and its supplementary materials.

