## Supplemental_Figure_1 for "Bulk sample testing by real-time PCR for detecting *Diaporthe destruens*-infected seedlings of sweet potatoes": Suppl Fig1 Probit regression results.pptx

### Slide 1
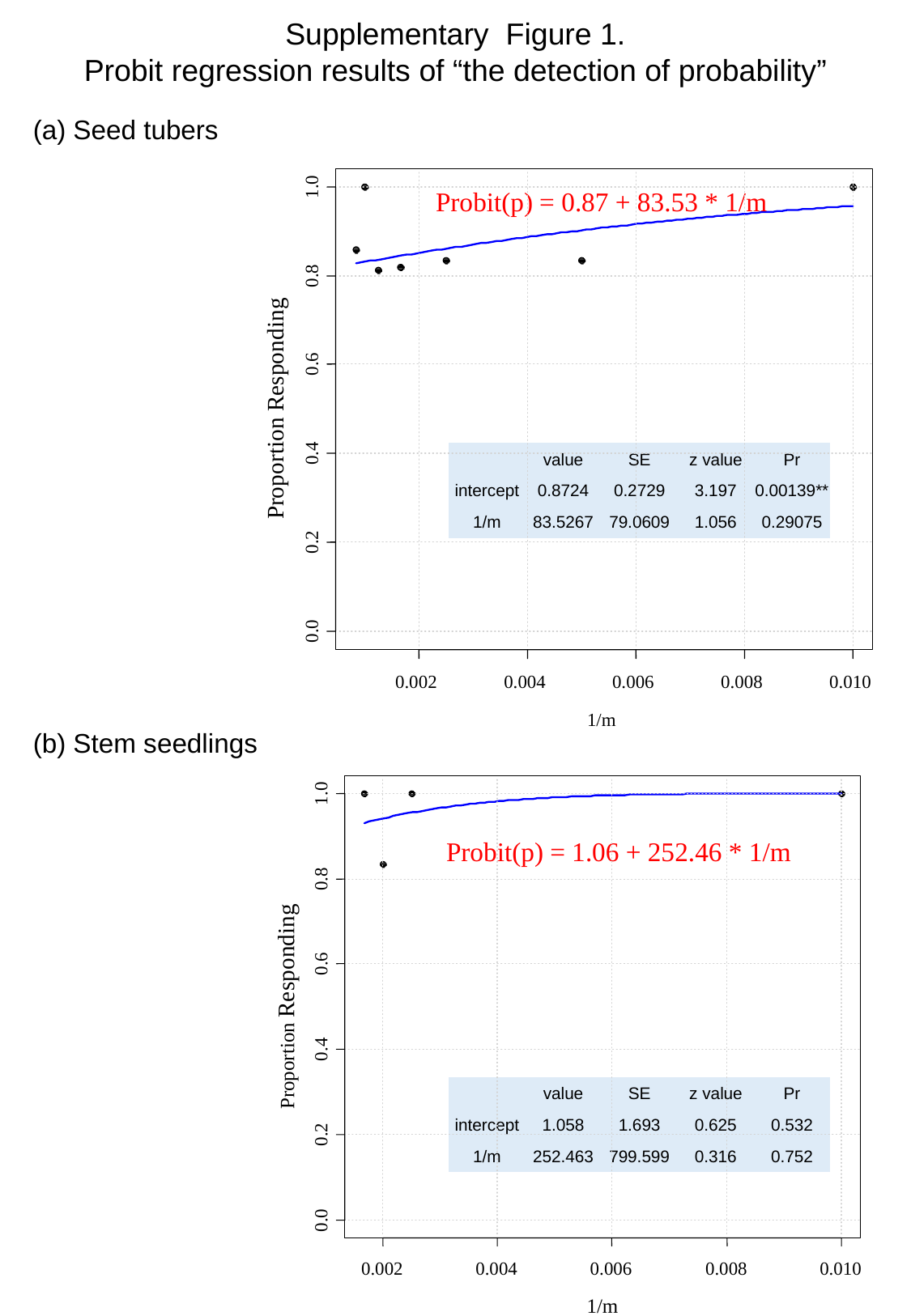

Supplementary Figure 1.
Probit regression results of “the detection of probability”
(a) Seed tubers
1.0
Probit(p) = 0.87 + 83.53 * 1/m
0.8
0.6
Proportion Responding
0.4
0.2
0.0
0.002
0.004
0.006
0.008
0.010
1/m
| | value | SE | z value | Pr |
| --- | --- | --- | --- | --- |
| intercept | 0.8724 | 0.2729 | 3.197 | 0.00139\*\* |
| 1/m | 83.5267 | 79.0609 | 1.056 | 0.29075 |
(b) Stem seedlings
1.0
Probit(p) = 1.06 + 252.46 * 1/m
0.8
0.6
Proportion Responding
0.4
0.2
0.0
0.002
0.004
0.006
0.008
0.010
1/m
| | value | SE | z value | Pr |
| --- | --- | --- | --- | --- |
| intercept | 1.058 | 1.693 | 0.625 | 0.532 |
| 1/m | 252.463 | 799.599 | 0.316 | 0.752 |
