## Supplementary_Text_1 for "Bulk sample testing by real-time PCR for detecting *Diaporthe destruens*-infected seedlings of sweet potatoes": Supplement Text 1. Probit regression code.docx

**Supplemental Text 1. Example of Probit regression generation code in R**

sink("motokusare.txt")

cat("

m r n

100 9 9

200 10 12

400 10 12

600 9 11

800 13 16

1000 7 7

1200 6 7

",fill=TRUE)

sink() # Data entry for multiple sample testing of seed tubers in Table 1A

motokusare <- read.table("motokusare.txt", header=TRUE)

motokusare$x <- 1/motokusare$m

motokusare$response <- cbind(motokusare$r,motokusare$n-motokusare$r)

\# Probit regression

motokusare.glm <- glm(response ~ x, family=binomial(link = "probit"), data = motokusare)

summary(motokusare.glm)

\# Estimated “the detection of probability (POD)” at 1/800 concentration

newdata <- data.frame(m=NA, r=NA, n=NA, x=0.00125, response=cbind(NA,NA)) #x=1/800=0.00125

motokusare.predict <- predict(motokusare.glm, newdata = newdata, type = "response")

\# POD at 1/800 concentration

motokusare.predict	# Results= 0.8356675

\# Non-POD at 1/800 concentration

1-motokusare.predict # Results= 0.1643325

<Display of probit regression curve>

\# Preparation of plot

\# Create a new data frame and calculate the predicted value for each x

new_data <- data.frame(x = seq(min(motokusare$x), max(motokusare$x), length.out = 100))

new_data$pred <- predict(motokusare.glm, newdata = new_data, type = "response")

\# Plotting actual data and predicted curves

plot(motokusare$x, motokusare$r / motokusare$n, xlab = "1/m", ylab = "Proportion Responding",

main = "Probit Regression", pch = 19, ylim = c(0, 1))

\# Draw the probit curve

lines(new_data$x, new_data$pred, col = "blue", lwd = 2)

\# Adding Grid Lines (optional)

grid()

\# Add a legend (optional)

legend("topright", legend = c("Observed Data", "Probit Model"), col = c("black", "blue"), pch = c(19, NA), lty = c(NA, 1), lwd = 2)

\# Get the model coefficients and create an approximate equation (optional)

coefficients <- coef(motokusare.glm)

intercept <- coefficients[1]

slope <- coefficients[2]

formula_text <- paste("Probit(y) = ", round(intercept, 3), " + ", round(slope, 3), " * 1/m")

\# Displaying approximate equations on a graph (optional)

text(x = min(motokusare$x), y = 0.9, labels = formula_text, pos = 4, col = "red")

cat("Probit(y) = ", intercept, " + ", slope, " * 1/m\n")
