## Supplemental_Text_2 for "Bulk sample testing by real-time PCR for detecting *Diaporthe destruens*-infected seedlings of sweet potatoes": Supplement Text 2. sampling number code.docx

**Supplemental Text 2. Example of Determining total sample size, number of subgroups, and number of samples within subgroups**

\# case of beta=0.05, p=0.01, phi=0.1875, when the total number n is divided into subgroups g of numbers from 1 to 10,

beta <- 0.05; p <- 0.01; phi <- 0.1875

for(g in 1:10){

if(beta^(1/g) > phi){

m <- ceiling(-log(1 - (1 - beta^(1/g))/(1-phi))/p); n<-g*m}

else {m <- NA; n <- NA};cat("g= ",g,", m= ",m,", n= ",n,"\n",sep="") }

\# case of beta=0,05, p=0.001, phi=0.1875, when the total number n is divided into subgroups g of numbers from 1 to 10,

beta <- 0.05; p <- 0.001; phi <- 0.1875

for(g in 1:10){

if(beta^(1/g) > phi){

m <- ceiling(-log(1 - (1 - beta^(1/g))/(1-phi))/p); n<-g*m}

else {m <- NA; n <- NA};cat("g= ",g,", m= ",m,", n= ",n,"\n",sep="") }
